# A decade of arbovirus emergence in the temperate southern cone of South America: dengue, *Aedes aegypti* and climate dynamics in Córdoba, Argentina

**DOI:** 10.1101/2020.01.16.908814

**Authors:** Elizabet L. Estallo, Rachel Sippy, Anna M. Stewart-Ibarra, Marta G. Grech, Elisabet M. Benitez, Francisco F. Ludueña-Almeida, Mariela Ainete, María Frias-Cespedes, Michael Robert, Moory M. Romero, Walter R. Almirón

## Abstract

**Background:** Argentina is located at the southern temperate range of arboviral transmission by the mosquito *Aedes aegypti* and has experienced a rapid increase in disease transmission in recent years. Here we present findings from an entomological surveillance study that began in Córdoba, Argentina, following the emergence of dengue in 2009.

**Methods:** From 2009 to 2017, larval surveys were conducted monthly, from November to May, in 600 randomly selected households distributed across the city. From 2009 to 2013, ovitraps (n=177) were sampled weekly to monitor the oviposition activity of *Ae. aegypti*. We explored seasonal and interannual dynamics of entomological variables and dengue transmission. Cross correlation analysis was used to identify significant lag periods.

**Results:** *Aedes aegypti* were detected over the entire study period, and abundance peaked during the summer months (January to March). We identified a considerable increase in the proportion of homes with juvenile *Ae. aegypti* over the study period (from 5.7% of homes in 2009-10 to 15.4% of homes in 2016-17). *Aedes aegypti* eggs per ovitrap and larval abundance were positively associated with temperature in the same month. Autochthonous dengue transmission peaked in April, following a peak in imported dengue cases in March; autochthonous dengue was not positively associated with vector or climate variables.

**Conclusions:** This longitudinal study provides insights into the complex dynamics of arbovirus transmission and vector populations in a temperate region of arbovirus emergence. Our findings suggest that Córdoba is well suited for arbovirus disease transmission, given the stable and abundant vector populations. Further studies are needed to better understand the role of regional human movement.

**Author summary:** There is an increasing risk of arbovirus transmission in temperate regions. Argentina is located at the southern range of dengue virus transmission by the *Aedes aegypti* mosquito. In the last decade, epidemics of dengue fever have emerged for the first time in the city of Córdoba, Argentina. We present the study design and findings from an entomological surveillance study in Córdoba, which began following the emergence of dengue in 2009. We found that *Ae. aegypti* were most abundant from January to March, followed by a peak in local dengue transmission in April. Over the study period, we noted a considerable increase in the proportion of homes with *Ae. aegypti*. Vector indices were positively associated with warmer temperatures, which have been increasing in this region. However, the timing of local dengue transmission appears to be driven by the appearance of imported dengue cases associated with human movement. These results highlight the important role of long term surveillance studies in areas of disease emergence.

## 1. Introduction

Over the last two decades, Argentina has experienced the emergence of epidemics of arboviral diseases vectored by *Aedes* mosquitoes (Aviles et al., 1999; Vezzani and Carbajo, 2008). Cases of dengue fever, chikungunya, and Zika have been reported from northern and central provinces (PAHO, 2019). Argentina is located at the southern limit of arboviral transmission by *Aedes aegypti*, the primary disease vector, thus the increasing burden of disease suggests a southern expansion of the vector and the arboviruses they carry (Masuh, 2008; Robert et al., 2019).

Córdoba, the second largest city in Argentina, experienced its first dengue outbreak in 2009 followed by subsequent outbreaks in 2013, 2015, and 2016 (Robert et al., 2019). In the midst of the COVID-19 global pandemic, Córdoba is currently experiencing the largest dengue outbreak to date (caused by DENV serotypes 1 and 4), which began in early 2020 (Ministerio de Salud de La Nación, 2020). Córdoba is among the southernmost cities in the Western Hemisphere to report autochthonous transmission of *Aedes*-borne arboviruses, highlighting its importance in the investigation of arbovirus emergence in the Americas and globally.

*Aedes aegypti* was considered eradicated from all of Argentina between 1963-1986 (Curto et al., 2002) and first re-appeared in Córdoba in 1996 (Almirón and Almeida, 1998). Since then, the vector has become established in the city. It is well adapted to the urban environment, living in and around human dwellings and feeding preferentially on human hosts (WHO, 2009).

Climate plays an important role in the *Ae. aegypti* life cycle as well as in its ability to transmit viruses. *Aedes aegypti* juvenile habitat consist of water-bearing containers such as man-made cisterns, discarded buckets, and tires (Manrique-Saide et al., 2011). Rainfall and drought can both potentially increase the availability of larval habitat, depending on local water storage practices and housing characteristics (Lowe et al., 2018; Stewart Ibarra et al., 2013). Prior studies in the subtropical northwest region of Argentina revealed a year-round vector population in areas with an annual mean temperature around 20°C. A low number of eggs was recorded during the winter season, and peak in oviposition was detected during summer months (December to March in the southern hemisphere) (Estallo et al., 2016, 2015, 2013; Micieli and Campos, 2003). Studies from the temperate central region, including Buenos Aires (Campos and Maciá, 1996; Carbajo et al., 2006, 2004; Micieli et al., 2006; Vezzani et al., 2004) and Córdoba provinces (Avilés et al., 1997; Domínguez et al., 2000; M. G. Grech, 2013), revealed that oviposition activity discontinued during the cool winter months due to low ambient temperatures (below 17°C).

Vector surveillance studies provide local empirical data on the distribution and density of arthropod species of public health concern. This information can be used to trigger public health alert systems, to guide decisions about interventions, and to evaluate control programs (Focks, 2003). These data can also be used to inform our understanding of the linkages between vector abundance, disease risk, and local climate conditions. In response to the first dengue outbreak in Córdoba in 2009, the province created the “Master Plan to Fight Dengue” and the Ministry of Health (MoH) implemented an ambitious *Ae. aegypti* surveillance program in cooperation with local academic partners.

Here we report on the study design, methods, and empirical data from this longitudinal surveillance study for the first time, following the first dengue outbreak in the temperate city of Córdoba. We describe the seasonal and annual variation in Ae. aegypti and dengue cases and explore the lagged associations with local climate variables. We explored the hypotheses that larval abundance is positively associated with warmer temperatures and greater rainfall in the months prior, and that dengue incidence is positively associated with greater vector abundance in the months prior. Few studies of this nature have been conducted in southern temperate zones of arbovirus emergence.

The data generated by this study lays the groundwork for further investigation regarding the dynamics of arbovirus emergence.

## 2. Methods

### 2.1 Ethics

Entomological data were collected from households by the MoH of Córdoba as part of the routine surveillance program, thus no ethical review or informed consent was required. Dengue case data at the city-level were extracted from National MoH bulletins and aggregated to weekly and monthly case counts; no identifying information was provided.

### 2.2 Study site

This study was conducted in the city of Córdoba (31.4°S, 64.2°W, population 1.3 million), which is located in the central region of Argentina (elevation: 360-480 m above sea level) (INDEC, 2010). The city has grown outward from the center to the periphery; agricultural fields with patches of forest surround the urban core (Maristany et al., 2008) (Fig. 1). The local climate of Córdoba is warm temperate with hot summers and four seasons (Cwa under the Köppen-Geiger classification system) (Peel et al., 2007). On average, the city center is several degrees warmer than the urban periphery, due to dense construction and location in a topographic depression (Asar et al., 2019). Summer is a warm, rainy season from November to March (monthly mean temp: 22.7°C, max temp: 37.6°C, min temp: 11.8°C, monthly total rainfall: 123.2 mm), and winter is a cool, dry season from June to September (monthly mean temp: 14.8°C, max temp: 30.9°C, min temp: 2.4°C, monthly total rainfall: 36.1 mm). *Aedes aegypti* is the only known vector of dengue in Córdoba. The vector does not reproduce (oviposit) during the winter, presumably due to cool temperatures (M. Grech, 2013); it is thought that the vectors persist through the cold season as egg. Vector control by the MoH is mostly focal around homes with dengue cases, and includes indoor and outdoor fumigation, larviciding with *Bacillus thuringiensis israelensis* (BTI), and eliminating standing water (Ministerio de Salud de la Nación, 2009).

**Fig 1.**
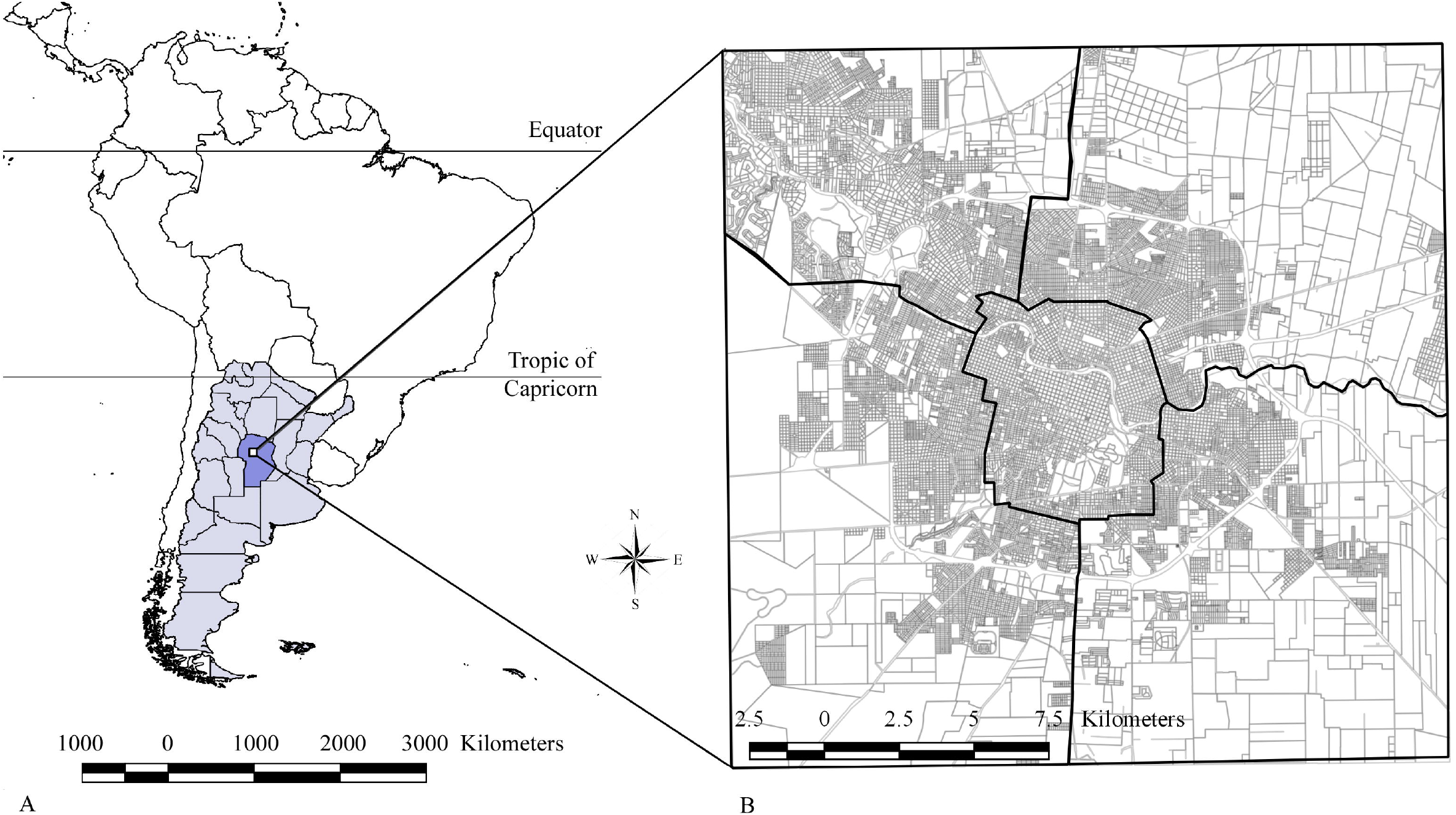
Location of Córdoba in South America. The location of Córdoba Province (dark purple) shown within the country of Argentina (light purple), with an inset map showing the city of Córdoba, the location of this study, with roads in grey and waterways in blue. Note that Córdoba is located in the temperate southern cone of South America.

Field studies were conducted under the *Ae. aegypti* surveillance program of the Department of Epidemiology of the Córdoba Province Ministry of Health in cooperation with the Córdoba Entomological Research Centre (CIEC) of Córdoba National University (UNC) and the Biological and Technological Research Institute (IIByT) of Córdoba National University (UNC) and the National Research Council (CONICET). No formal ethical review or consent procedure was required for this study, as the field operations were conducted as part of routine surveillance activities by the MoH. Entomological samples were collected from households that verbally agreed to participate in the study and were taken to the CIEC laboratory to be processed.

### 2.3 Larval sampling

From 2009 to 2017, the distribution and abundance of *Ae. aegypti* were monitored through monthly larval surveys conducted across the city. Sample periods each year were chosen by the CIEC laboratory team to span the period before, during and after mosquito activity peak in the region (Domínguez et al., 2000). Each month, 600 different homes were randomly selected for sampling. The city was divided into 215 quadrants (1.2 km per side), which were distributed across five areas of approximately the same land area: Central, Northeast, Northwest, Southeast, and Southwest (Fig 1). In each quadrant, we randomly selected 6 neighborhoods, and the field technicians randomly selected 20 homes to be inspected per neighborhood.

At each home, an inspector noted all containers inside and outside the home with standing water. Inspectors noted the presence of juvenile mosquitoes and collected specimens for species identification. Whenever possible, all juvenile mosquitoes were collected; for large containers where it was not possible to collect all specimens, an inspector collected three samples using a white dipper (62 ml volume). Specimens were transported to the CIEC laboratory. Pupae were reared to adults, and 3^rd^ and 4^th^ instar larvae were preserved in 80% ethanol and were identified using taxonomic keys (Darsie Jr, 1985). First and 2^nd^ instar larvae were reared until reaching the 3^rd^ instar in plastic trays with 500 ml of water from the natural larval habitat or dichlorine water. Each day larvae were fed 0.25 mg of liver powder per larva, and we cleaned the surface of the water using absorbent paper to avoid contamination by fungi and/or bacteria. The juvenile mosquitoes were counted and identified to species. For the purposes of this study, data were aggregated to the city level and we calculated the proportion of households and neighborhoods with juvenile *Ae. aegypti* present during each sampling period.

### 2.4 Ovitraps

The weekly oviposition activity of *Ae. aegypti* was observed from November to May over four years (2009 – 2013) using ovitraps, a sensitive method of detecting the presence of gravid female of *Ae. aegypti* (Fay & Eliason, 1966). Ovitraps were placed in randomly selected households (N=177) that were distributed evenly across the city. To assess oviposition during the winter months, a subset of evenly distributed ovitraps (n=40) were selected to continue sampling during June (late autumn-early austral winter seasons) to September 2010 (late winter-early austral spring seasons).

Ovitraps consisted of plastic bottles (350 ml volume, 8 cm diameter x 13 cm height) with filter paper as an oviposition substrate. An attractive infusion (250 ml) was prepared by fermenting dry cut grass with tap water (Reiter et al., 1991). Each week the traps were inspected and replaced in the same dwelling during the sampling years. Traps were transferred to the laboratory and the number of *Ae. aegypti* eggs per trap were counted using a stereomicroscope. For the purposes of this study, ovitrap data were aggregated to the city level. We calculated the proportion of traps that were positive/negative for *Ae. aegypti* eggs and the mean number of eggs per trap during each sampling period.

### 2.5 Climate information

The Argentinian National Meteorological Service provided daily weather data for the study period (rainfall, min/mean/max temperature, min/mean/max relative humidity) from the Observatory station (31.42°S; 64.20°W). This is one of the oldest weather stations in Latin America (in operation since 1871) and the most reliable in Córdoba (de la Casa and Nasello, 2010). We calculated summary monthly values (mean, standard deviation) over the study period, summary values by epidemiologic week, and summary values corresponding with the timing of ovitrap and larval survey collection periods.

### 2.6 Dengue case data

Weekly dengue case reports (January 2009 to December 2017) were extracted from the weekly epidemiological bulletins of the Argentina Health Secretary (Ministerio de Salud de Nación (MSN)., 2017). Dengue diagnostic procedures, case definitions and data extraction procedures have been previously described (Robert et al., 2019). Cases include suspected and laboratory confirmed cases aggregated at the city-level and annual incidence was calculated using the total city population for the corresponding year. Autochthonous and imported cases are reported separately.

### 2.7 Statistical analysis

Statistical analyses were conducted using time series of vector variables, dengue incidence (Supplemental Fig 1), and local climate (Supplemental Fig 2) at the city-level using R version 3.3.3 (*R Studio*, 2015) using the packages splines, TSA, geepack, MASS, lubridate, TSimpute (Bates et al., 2011; Chan and Ripley, 2012; Grolemund and Wickham, 2011; Halekoh et al., 2006; Moritz and Bartz-Beielstein, 2017; Team, 2018; Venables and Ripley, 2013). Missing values in the ovitrap data (n=10) were imputed via seasonally decomposed structural modeling with Kalman smoothing. Missing values in the larval data (n=20) were imputed via seasonally decomposed random selection. For each climate, mosquito abundance, and dengue incidence variable, we performed a spectral analysis and subsequently tested for the presence of inter-annual variability with a restricted cubic spline using model fit to determine the periodic frequency and the number of knots. These descriptive analyses formally test for the presence of periodicity in a time series and can be used to determine frequency and timing of peaks in the time series. Assessing inter-annual variability can detect the presence of trends in data covering multiple years. Models were fit using a generalized linear model with appropriate distributions for each variable using generalized estimating equations (auto-regressive correlation); best fit was determined using quasi-likelihood information criteria (QIC) as compared to a null model. Residual plots and model assumptions were examined to inform final model selection. Data were analyzed in relation to the collection periods of each respective variable; means and frequencies for these collection periods were calculated for reference.

We assumed a unidirectional temporal relationship between the variables as follows: climate affecting all other variables, *Ae. aegypti* eggs affecting *Ae. aegypti* larvae and dengue incidence, and *Ae. aegypti* larvae variables affecting dengue incidence. Both autochthonous and imported cases were included in the analysis. To examine the correlations between these variables, cross-correlation functions were calculated between differenced monthly summary data for each variable with lags up to two months. We selected this period as it is a biologically plausible period of time that includes the combined time for *Ae. aegypti* egg hatching and larval development to adult mosquitoes.

## 3. Results

### 3.1 *Aedes aegypti* detection

*Aedes aegypti* were detected over the entire study period, with the exception of the winter months (June to September), where winter was sampled in 2010 only. There was a notable increase in *Ae. aegypti* larval abundance over the study period (Supplemental Fig. 1). On average, *Ae. aegypti* abundance peaked once annually (from January to March), followed by a peak in autochthonous dengue transmission in April (Supplemental Fig 1). Our findings support the hypothesis that larval abundance was associated with warmer temperatures in months prior; however, autochthonous dengue transmission was not positively associated with larval abundance or climate variables. Results are described in the following and summary statistics for all variables are shown in S1 Table.

*Aedes aegypti* egg abundance was highest in January (Fig. 2), with a median of 33 eggs per trap and 61% of traps positive for eggs during this month. *Aedes aegypti* egg abundance was highest in the 2009-10 and 2011-12 seasons (Fig. 2), with a mean of 21 eggs per ovitrap per month for both seasons, and 36.4% and 41.6% of traps positive for *Ae. aegypti* eggs in 2009-10 and 2011-12, respectively. Note that ovitraps were used only until 2013 (4 collection seasons).

**Fig 2.**
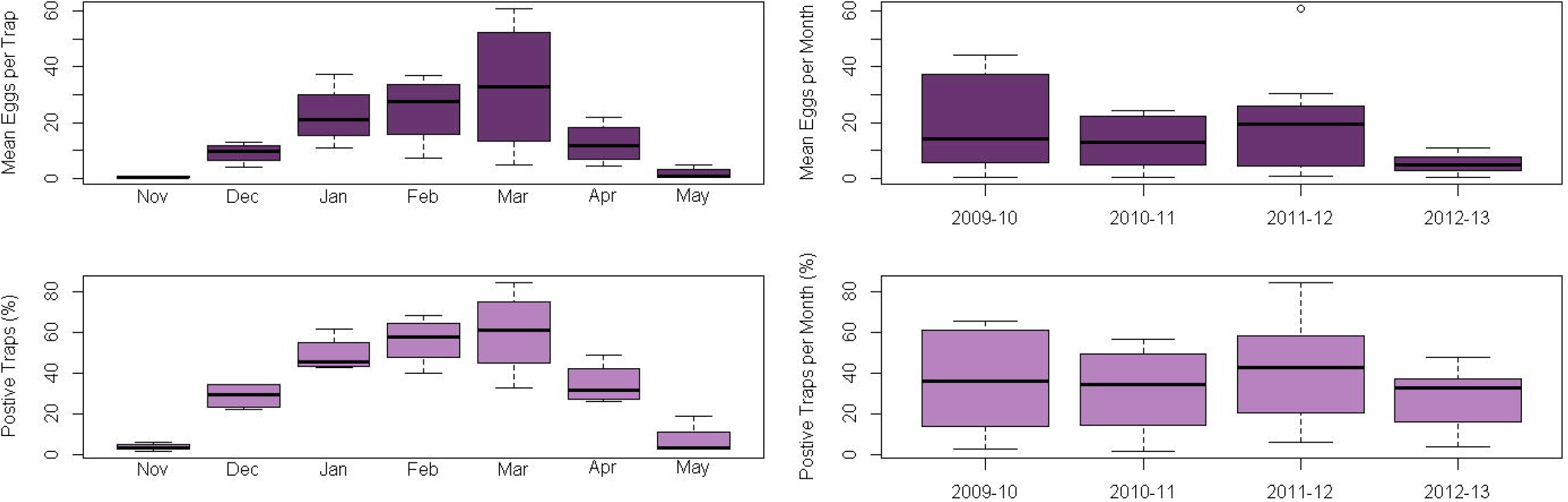
Seasonal and annual *Aedes aegypti* egg counts and positivity from ovitraps (2009-2013). Box and whisker plots show the monthly (left) and annual (right) median and quartiles of *Ae. aegypti* eggs counts and positivity of ovitraps distributed across the city of Córdoba from 2009 to 2013. Note that ovitraps were sampled during the winter months (June to September) only in 2010. Annual averages calculated using data collected during the sampling season (November—May) each year, from 2009 to 2013. Top: Number of *Ae. aegypti* eggs collected per ovitrap (left) or per ovitrap per annual sampling season (right). Bottom: The percent of ovitraps with *Ae. aegypti* eggs (left) or per annual sampling season (right).

*Aedes aegypti* larval abundance was highest in March (at the residences level) and February (at the neighborhood level) in each year (Fig. 3). During these months, 14% of residences and 85% of neighborhoods had containers positive for larvae. Larval abundance increased over the study period, with the highest infestation levels detected in 2015-16 and 2016-17 (Fig. 3). Fourteen percent and 15.8% of residences, and 83.1 and 71.3% of neighborhoods, had containers positive for *Ae. aegypti* larvae in 2015-16 and 2016-17, respectively.

**Fig 3.**
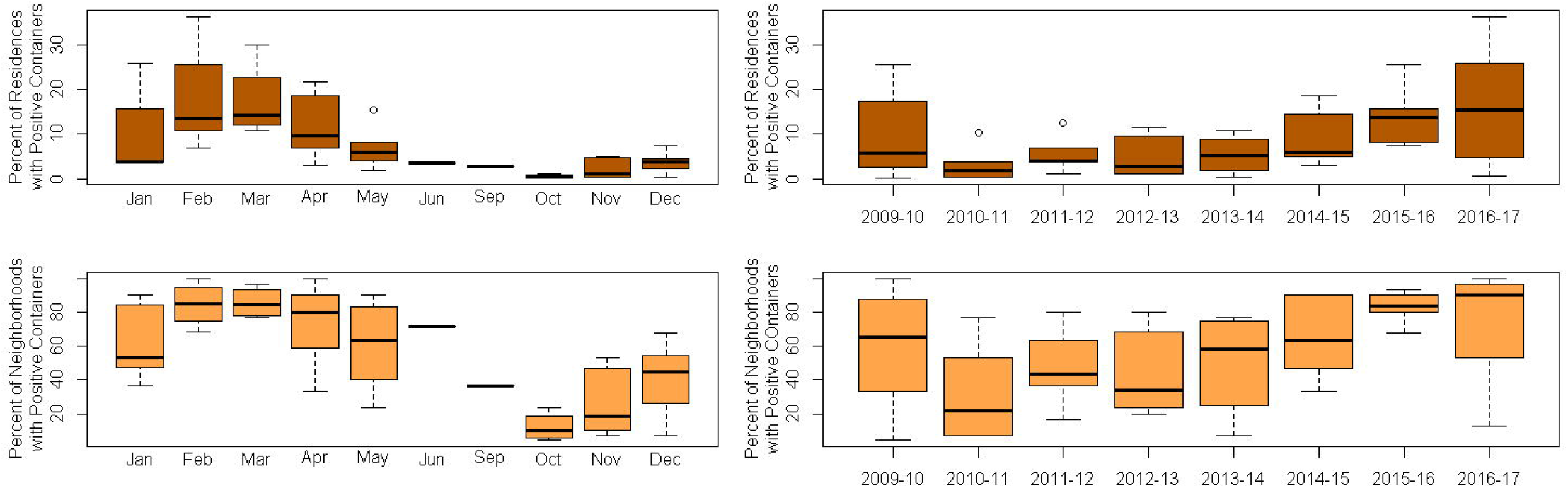
Seasonal and annual *Aedes aegypti* larval abundance from household larval surveys (2009-2017). Box and whisker plots show the monthly (left) and annual (right) median and quartiles of *Ae. aegypti* larval abundance from household larval surveys conducted in households in Córdoba from 2009 to 2017. Annual averages were calculated using data collected during the sampling season each year (November—May). Top: Percent of homes with water-bearing containers with juvenile *Ae. aegypti*. Bottom: Percent of neighborhoods with water-bearing containers with juvenile *Ae. aegypti*.

Dengue cases were highest during April across all years, with a local transmission season from March to May (mean 65 autochthonous cases and mean 9 imported cases, Fig. 4). The number of dengue cases was highest in 2016 (Fig. 4), with 689 total autochthonous and 139 total imported cases.

**Fig 4.**
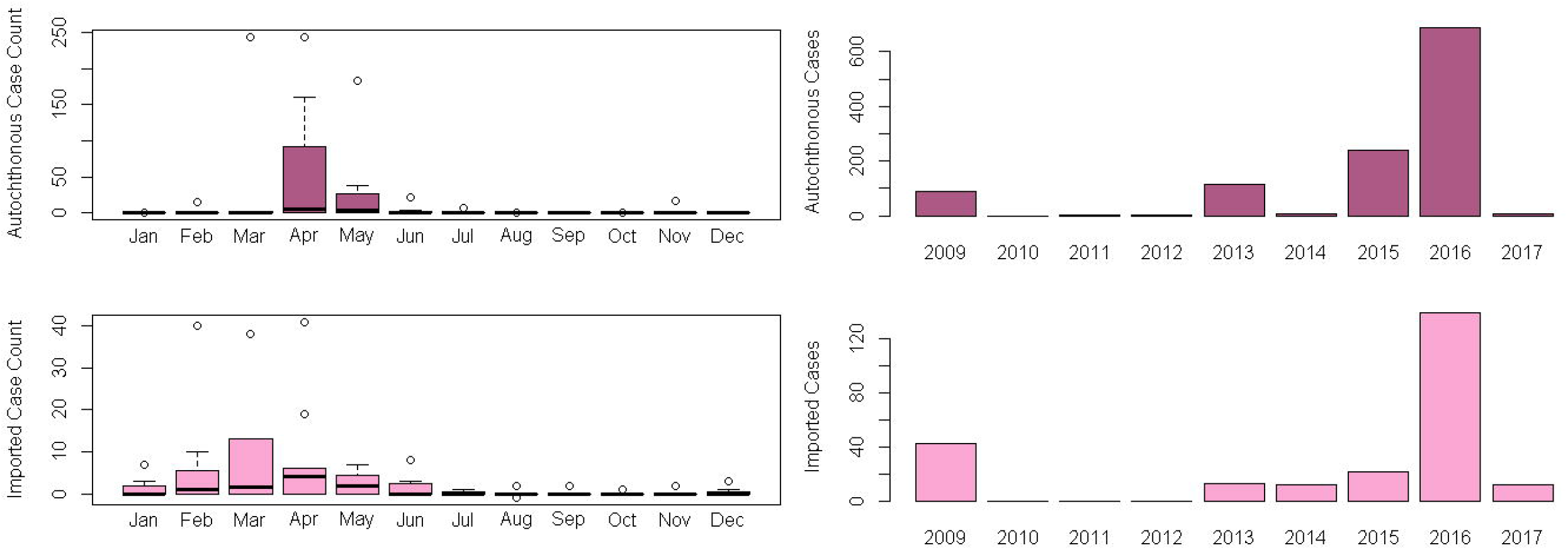
Seasonal and annual imported and autochthonous dengue cases in the city of Córdoba (2009-2017). Box and whisker plots (left) show the monthly median and quartiles of dengue cases reported in Córdoba from 2009 to 2017. Annual (right) imported and autochthonous dengue cases in Córdoba as reported by the Ministry of Health from 2009 to 2017. Top: Total annual reported autochthonous dengue cases (no travel history). Bottom: Total annual imported dengue cases.

### 3.2 Periodicity

The results of the seasonal analysis for climate, mosquito abundance, and dengue incidence variables are presented in Supplemental Table 2. All climate variables were found to have a 12-month periodicity with nonlinear interannual variability, except for minimum relative humidity, which had no interannual variability, and maximum relative humidity, which had no periodicity nor interannual variability (*i.e*. temporal trend). The detection of temporal trends can be limited with shorter sets of time series data.

*Aedes aegypti* eggs were collected from ovitraps every 7 days on average; an average of 26 times within each collection season. Both *Ae. aegypti* egg abundance variables (trap positivity and mean number of eggs) exhibited a peak about once each year on average, and the mean number of eggs had an additional peak occurring every 2 years on average. Trap positivity had no interannual variability while the mean number of eggs had nonlinear interannual variability.

Over the study period, on average, *Ae. aegypti* larvae were collected every 39 days, six times per collection season. *Aedes aegypti* larvae abundance variables (proportion of positive homes, proportion of positive neighborhoods) were found to have different seasonal patterns: there was no periodicity nor interannual variability for the proportion of positive homes, but the proportion of positive neighborhoods peaked once per year, as well as every 10 years.

Dengue surveillance reports of autochthonous and imported cases occurred every 8 days on average; an average of 39 times per year. Autochthonous dengue incidence peaked approximately twice per year and annually. Imported dengue incidence peaked approximately every 1.5 and 3 years. There was no interannual variability in autochthonous or imported dengue incidence.

### 3.3 Cross correlation analysis

The results of the cross-correlation between *Ae. aegypti* eggs and larval abundance, climate, and dengue incidence variables are shown in Table 1. Cross-correlation function plots for all tested relationships are available in Supplemental Fig. 3 *Ae. aegypti* egg abundance (mean number of eggs per ovitrap) was positively correlated with mean temperature in the same month. *Aedes aegypti* larval abundance (percent of neighborhoods with positive larval traps) was positively correlated with minimum temperature in the same month, while the percentage of residences with positive larval traps was not correlated with any climate variable. *Aedes aegypti* egg abundance was positively correlated with larval abundance (percent of neighborhoods with positive larval traps) at lag 1.

**Table 1.**
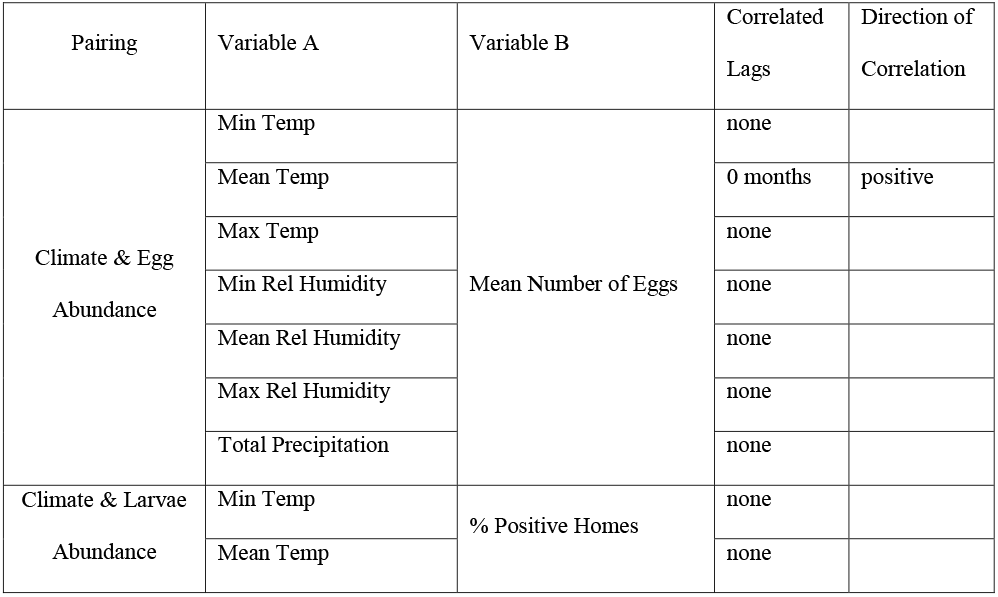
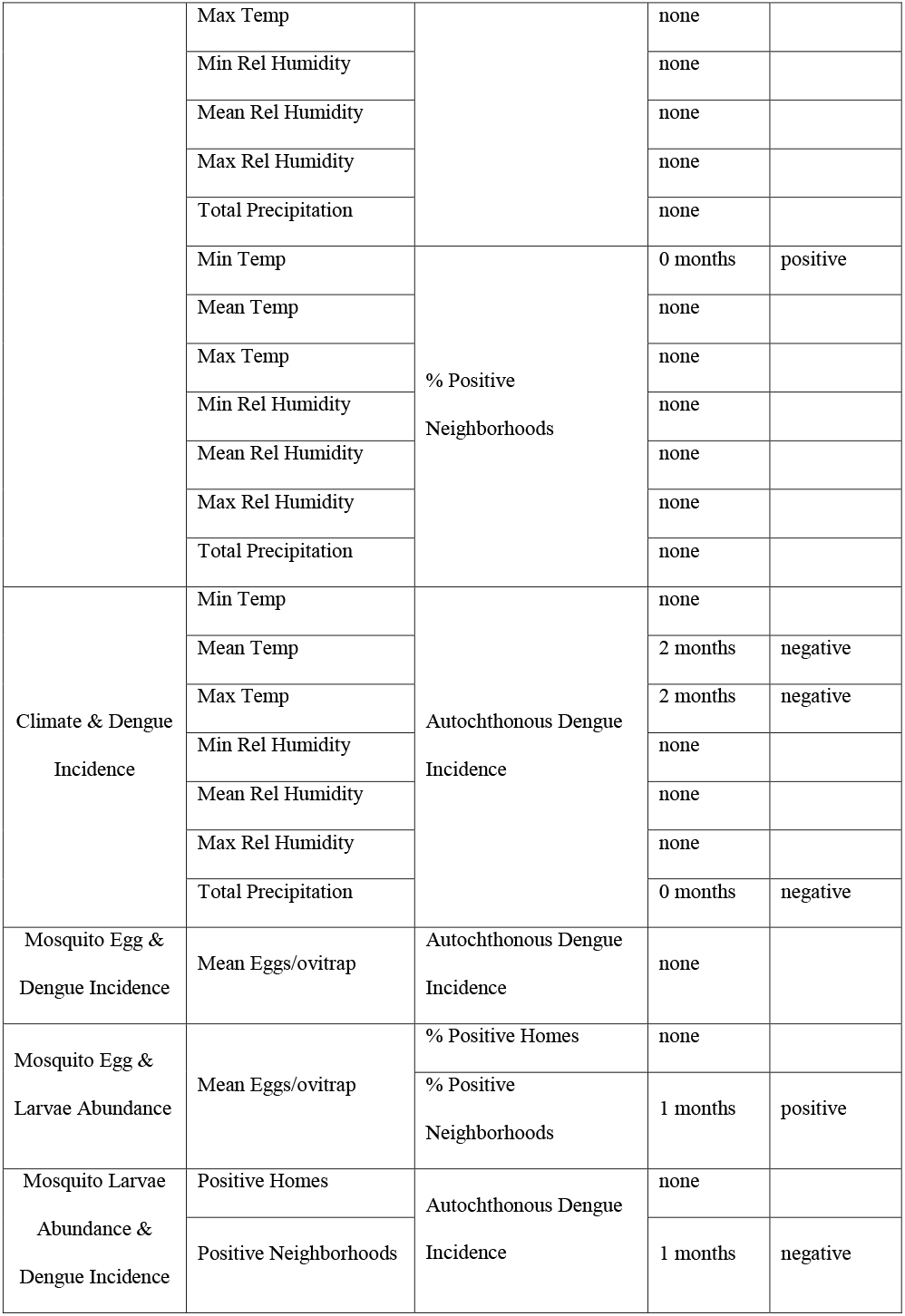
Cross-Correlation Between Climate, Mosquito Abundance, and Dengue Cases for 2009—2017. Using the cross-correlation function, we examined the correlation between pairs of variables, with variable A hypothesized to occur 0 to 2 months before variable B.

Autochthonous dengue incidence was negatively correlated with mean and maximum temperature at lag 2, and negatively correlated with total precipitation in the same month, with no correlation to any relative humidity variables. Autochthonous dengue incidence was not correlated with *Ae. aegypti* egg abundance. *Aedes aegypti* larvae abundance (percent of neighborhoods with larval presence) was negatively correlated with autochthonous dengue incidence at lag 1.

## 4. Discussion

Herein we present the findings from a vector surveillance study in the temperate city of Córdoba, Argentina, during the decade after which the first major epidemic of dengue occurred in the city. This study is among few such studies investigating arbovirus emergence in a temperate climate as it is actively occurring, providing a unique dataset with insights into dengue emergence and the drivers thereof. Among the key findings in this work, we identified a notable increase in the proportion of homes with juvenile *Ae. aegypti* (from 5.7% of homes in 2009-10 to 15.4% of homes in 2016-17). We also found that vector indices increased with warmer temperatures. In prior research, we noted an increase in ambient temperature anomalies over the last 30 years in Córdoba, suggesting a warming climate that could increase suitability for the mosquito vector (Robert et al., 2020). We did not identify positive associations between dengue disease transmission and lagged climate or vector variables, suggesting that disease dynamics in this zone of emergence may be driven more by intrinsic disease dynamics (e.g., herd immunity, importation of new serotypes) than by external environmental factors.

Prior research has shown that temperature is a critical driver of *Ae. aegypti* dynamics due largely to temperature effects on mosquito physiology (Mordecai et al., 2019). In our investigation, we found that the temperatures of Córdoba were warm enough to support the survival and breeding of *Ae. aegypti* populations from October to May. *Aedes aegypti* abundance peaked once annually (from January to March), followed by a peak in autochthonous dengue cases in April. Prior studies from Córdoba found that *Ae. aegypti* egg-to-adult survival began at mean minimum ambient temperatures greater than 13°C (Domínguez et al., 2000), suggesting that *Ae. aegypti* populations in Córdoba are not likely to survive when temperatures drop below this threshold. Taken together, these findings indicate that the temporal window for a stable *Ae. aegypti* population in Córdoba is rather short, which limits the period in which potential autochthonous dengue can occur. Indeed, Estallo *et al*. (2014) showed that low autumnal temperatures were an important factor limiting the spread of dengue during the first dengue outbreak in 2009.

Of note, we found that imported cases had an annual seasonal peak in March, followed by a peak in autochthonous transmission in April. Local dengue transmission was not positively associated with lagged climate or vector variables. We suspect that the negative associations with lagged climate variables may be spurious, and likely due to the timing of seasonal human movement to neighboring endemic regions within and outside the country (Estallo et al., 2014). For example, during 2009 dengue outbreak in the city, the introduction of DENV was associated with outbreaks in the neighboring countries of Bolivia, Brazil and Paraguay at the end of 2008, and also with dengue transmission in the Northern provinces of Argentina, where 92% of dengue cases in the country occurred that year (Estallo et al., 2014). It is possible that human movement into Córdoba from neighboring areas corresponds with summer breaks in academic schedules in Argentina (December-February) before university classes begin each year in March. An estimated 60% of the approximately 132,000 university students in the National University of Córdoba originate from other provinces or countries (Telediario Digital, 2014; Universidad Nacional de Cordoba, 2017), which could lead to importation of cases by students who travel between Córdoba and other dengue-endemic regions during the summer months. A prior study in Brazil also found that dengue transmission was most closely related to the movement of viremic humans rather than vector densities (Honório et al., 2009).

This study contributes to a growing body of knowledge on the linkages among vector abundance, disease cases, and climate. Local studies such as this are necessary, as these linkages may vary in space and time when comparing regions with distinct eco-epidemiological contexts (e.g., tropical endemic region versus temperate emergence region) and when different entomological surveillance methodologies are used (e.g., ovitraps, larval surveys, adult trapping). The lack of association between dengue and juvenile *Ae. aegypti* abundance was confirmed in prior studies in Córdoba (Estallo et al., 2014). Studies in Venezuela and Malaysia similarly found no relation (Barrera et al., 2002; Sulaiman et al., 1996), whereas studies in Puerto Rico, Vietnam, and Trinidad found that vector density measurements were associated with dengue transmission (Barrera et al., 2011; Chadee, 2009; Pham et al., 2011). It is possible that surveillance of juvenile vectors did not capture the entomological risk presented by adults of *Ae. aegypti* and that spatial variation in entomological risk leads to localized hotspots of transmission risk that are not captured in the aggregated city-level data. We also hypothesize that in a zone of emergence, the timing of transmission depends on the timing of the introduction of the virus (as a high proportion of the population is immunologically naïve to dengue) and transmission may be sustained with low vector densities.

As indicated by the appearance and continued transmission of dengue in Córdoba, this temperate region is actively experiencing the emergence of dengue (Masuh, 2008; Robert et al., 2019). The ongoing dengue outbreak is the largest to date, with over 4,000 cases reported thus far (Ministerio de Salud de La Nación, 2020). Emergence of dengue in temperate regions has been occurring with more frequency in the past two decades (Lourenço & Recker, 2014; Messina *et al*., 2014; Radke *et al*., 2012). As the climate becomes increasingly favorable for *Ae. aegypti* in Argentina, we anticipate that the southern range of the vector will continue to increase, with prolonged periods of activity during the year. Empirical studies such as this, from the lower thermal range of arbovirus transmission, can be used to improve models that investigate arbovirus dynamics in temperate zones of emergence, predict future outbreaks, and explore potential control measures.

It is important to note that we do not expect that the reported correlations between climate, mosquito abundance, dengue incidence to be causal in any way. Cross-correlation analyses do not adjust for potential confounding and we expect that the correlation found is due to the coincidental timing of Argentina vacation periods and peak larval activity. One limitation of our work is irregular sampling in the surveillance data, creating gaps in some periods, though the sampling design was strong and carefully designed by the team of researchers at the CIEC. It is also worth noting that the monthly summaries used in this study do not allow for an analysis of finer scale climate effects on mosquito vectors (e.g., daily diurnal temperature fluctuations) (Lambrechts et al., 2011). In addition, the shorter time series of ovitrap data does not allow for a more robust comparison to dengue transmission data.

## 5. Conclusions

This longitudinal study provides insights into the complex dynamics of arbovirus transmission and vector populations in a temperate region of arbovirus emergence. Our findings suggest that Córdoba is well suited for arbovirus disease transmission, given the stable and abundant vector populations. Further studies are needed to better understand the role of regional human movement, including origins, destinations, and timing, throughout South America to understand local patterns of disease transmission.

## Supporting information

Time series of monthly dengue, Aedes aegypti larval and Ae. aegypti eggs abundance.

Times series of monthly climate variables.

## Acknowledgements

The authors wish to acknowledge the United States Embassy in Argentina and Fulbright Commission as well as the Department of Epidemiology of the Córdoba Province Ministry of Health. AMSI and MAR were support in Córdoba Argentina by the USA Zika program of the United States Embassy in Argentina administered by the Fulbright commission. ELE, MGG, and WRA are members of the Consejo de Investigaciones Cientificas y Tecnológicas (CONICET) from Argentina, EMB is a PhD Student with scholarship support from CONICET.

---

**Supplemental Figure 1. Time series of monthly dengue, *Aedes aegypti* larval and *Ae. aegypti* eggs abundance.** Top: Dengue cases. Center: proportion of homes with *Ae. aegypti* larvas. Bottom: proportion of ovitraps (N=177) positive for *Ae. aegypti* and the mean number of eggs per ovitrap.

**Supplemental Figure 2. Times series of monthly climate variables.** Mean monthly temperatures (°Celsius) are in the top panel (red) (minimum and maximum in dashed lines), total monthly precipitation (mm) is in the central panel (blue) and mean monthly relative humidity (percentage) is in the bottom panel (green) (minimum and maximum in dashed lines).

**Supplemental Figure 3. Cross-correlation function plots for climate variables, *Aedes aegypti* abundance, and dengue cases.** Cross-correlation function (CCF) plots are given for tested correlations between climate variables and egg abundance (a), climate variables and larval abundance (b), climate variables and dengue incidence (c), egg abundance and dengue incidence (d), egg abundance and larval abundance (e), and larval abundance and dengue incidence (f). We assumed a unidirectional temporal relationship between the variables as follows: climate affecting all other variables, *Aedes aegypti* eggs affecting *Aedes aegypti* larvae and dengue incidence, and *Aedes aegypti* larvae variables affecting dengue incidence. Lags of 0-2 months were tested.

**Supplemental Table 1. Summary Statistics of Climate, *Aedes aegypti*, and Dengue Measures.** Mean and range by year are given for climate, *Aedes aegypti*, and dengue measures used in this study.

**Supplemental Table 2. Seasonality and Interannual Variability of Data.**

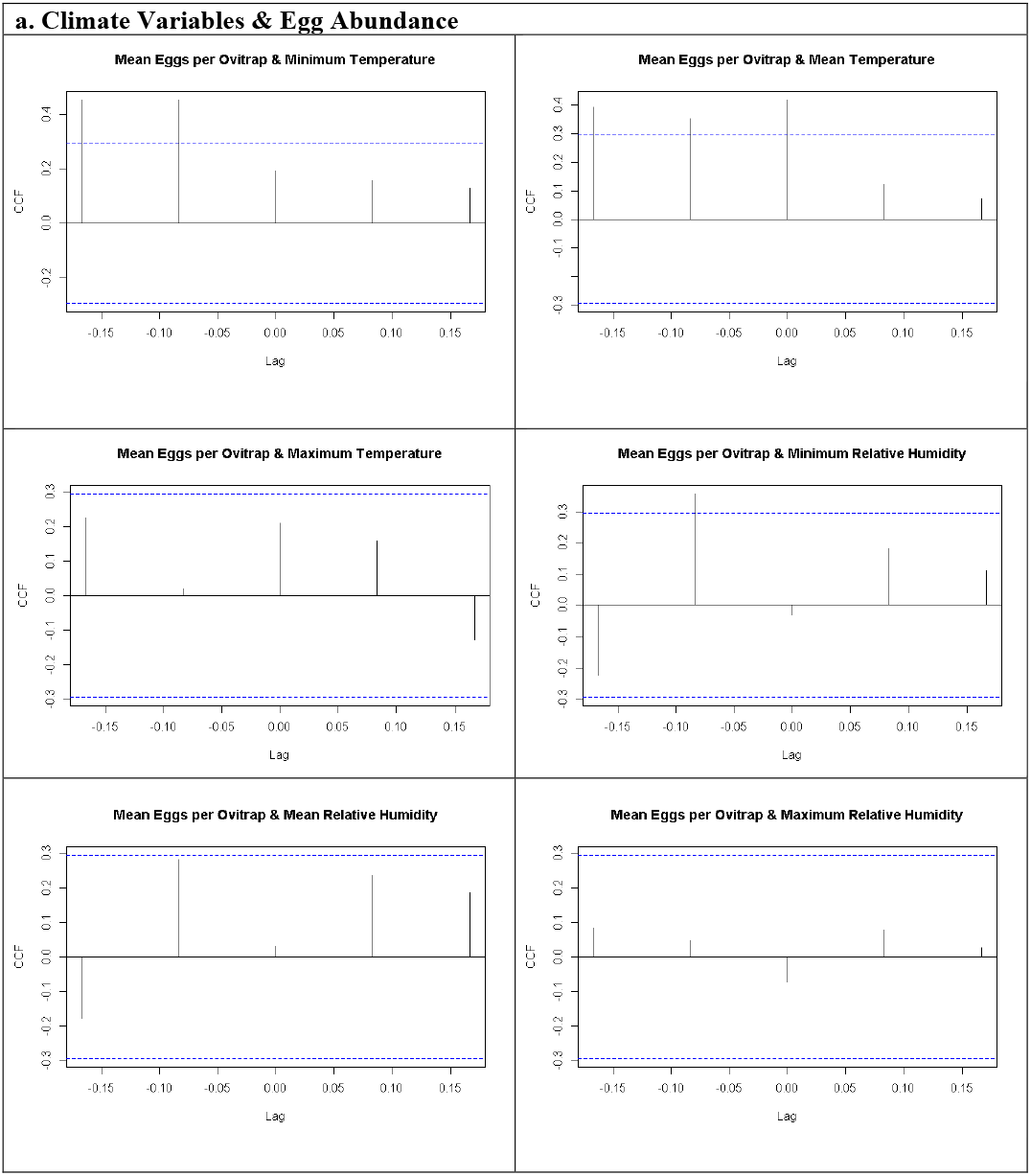
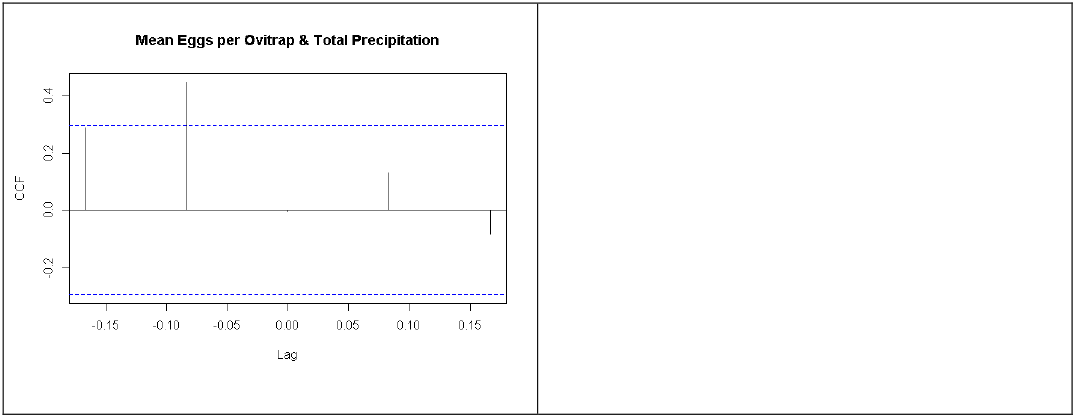
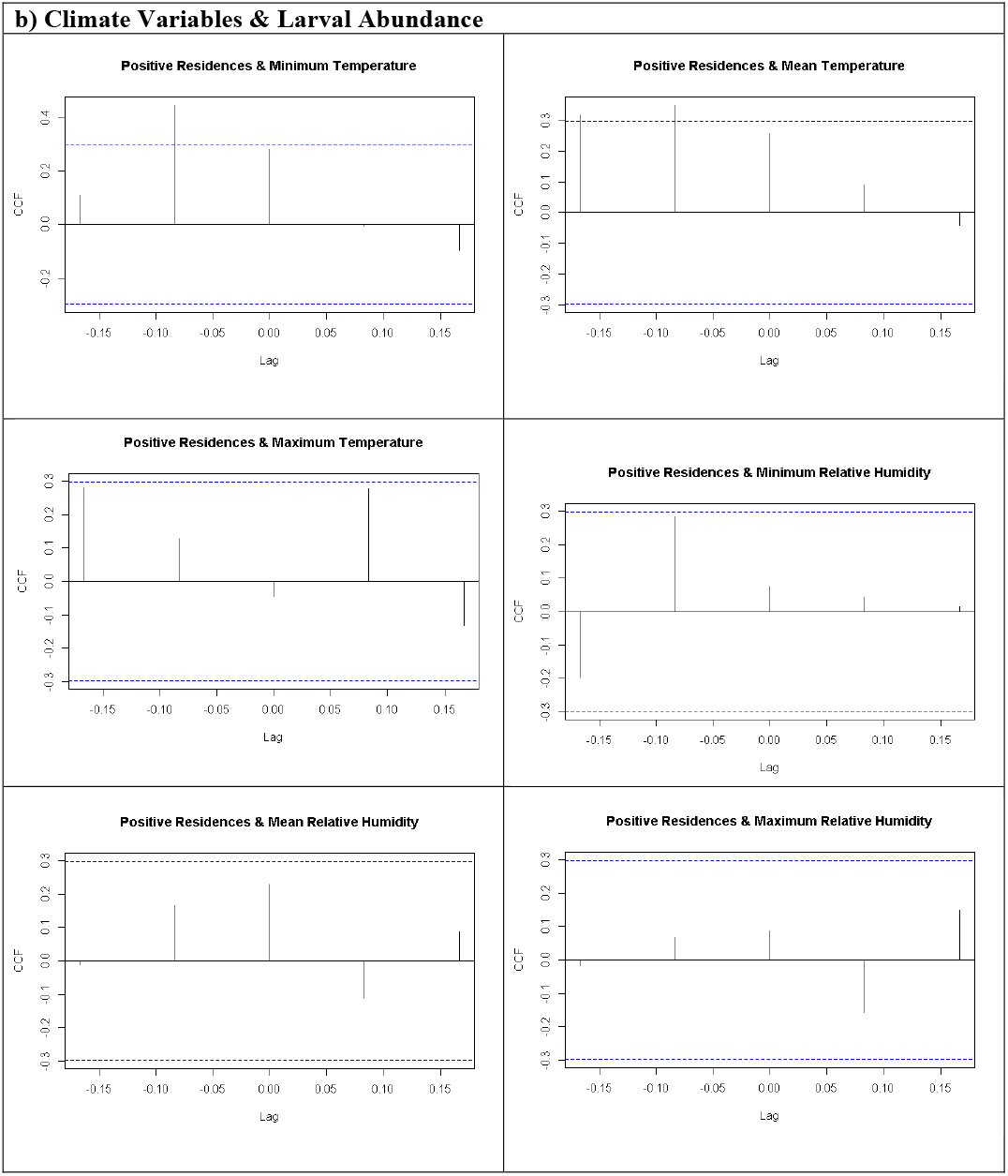
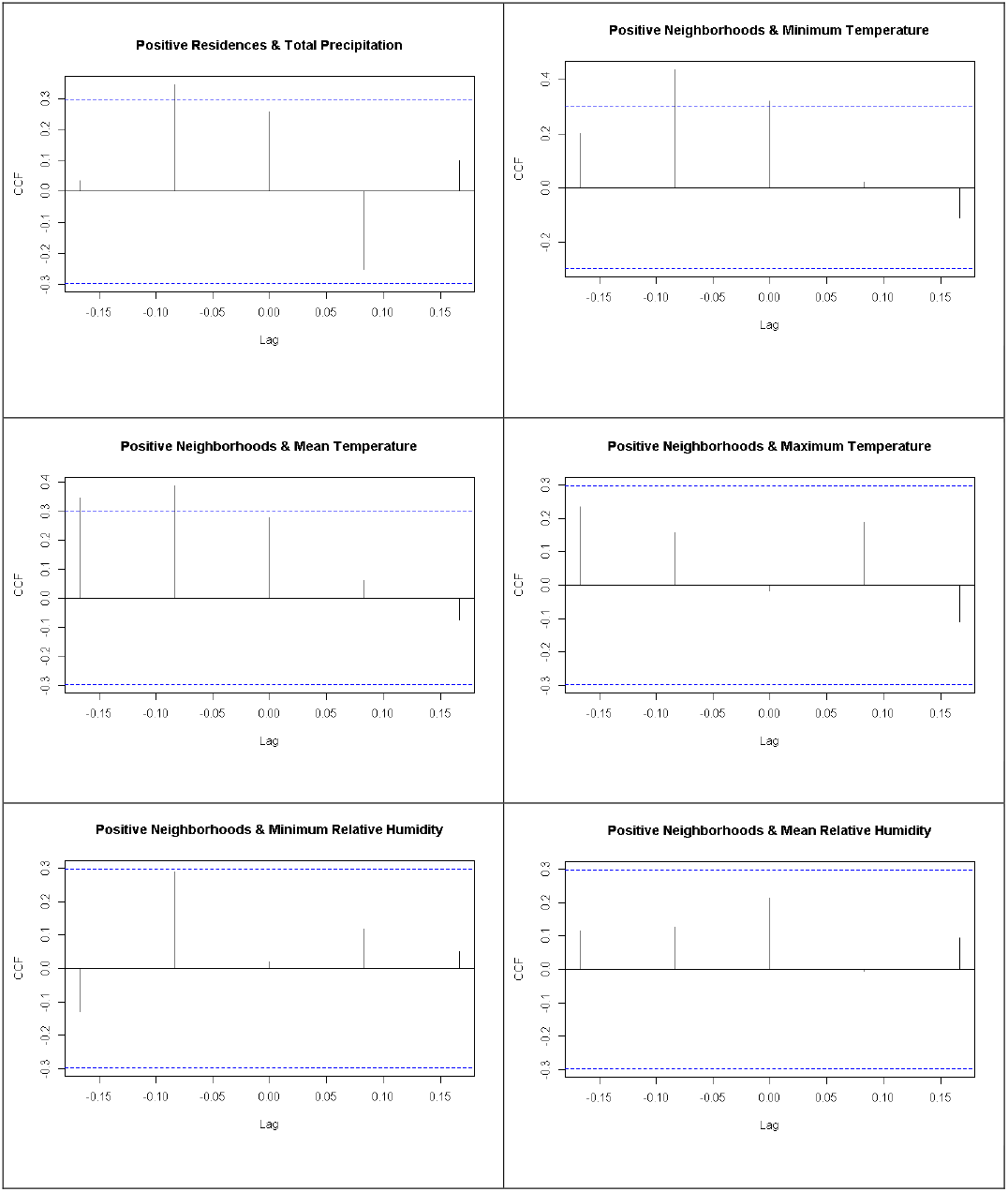
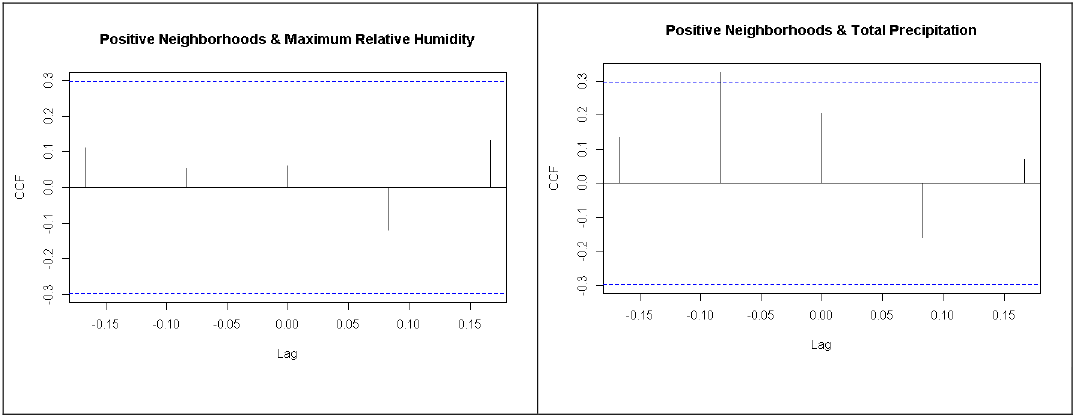
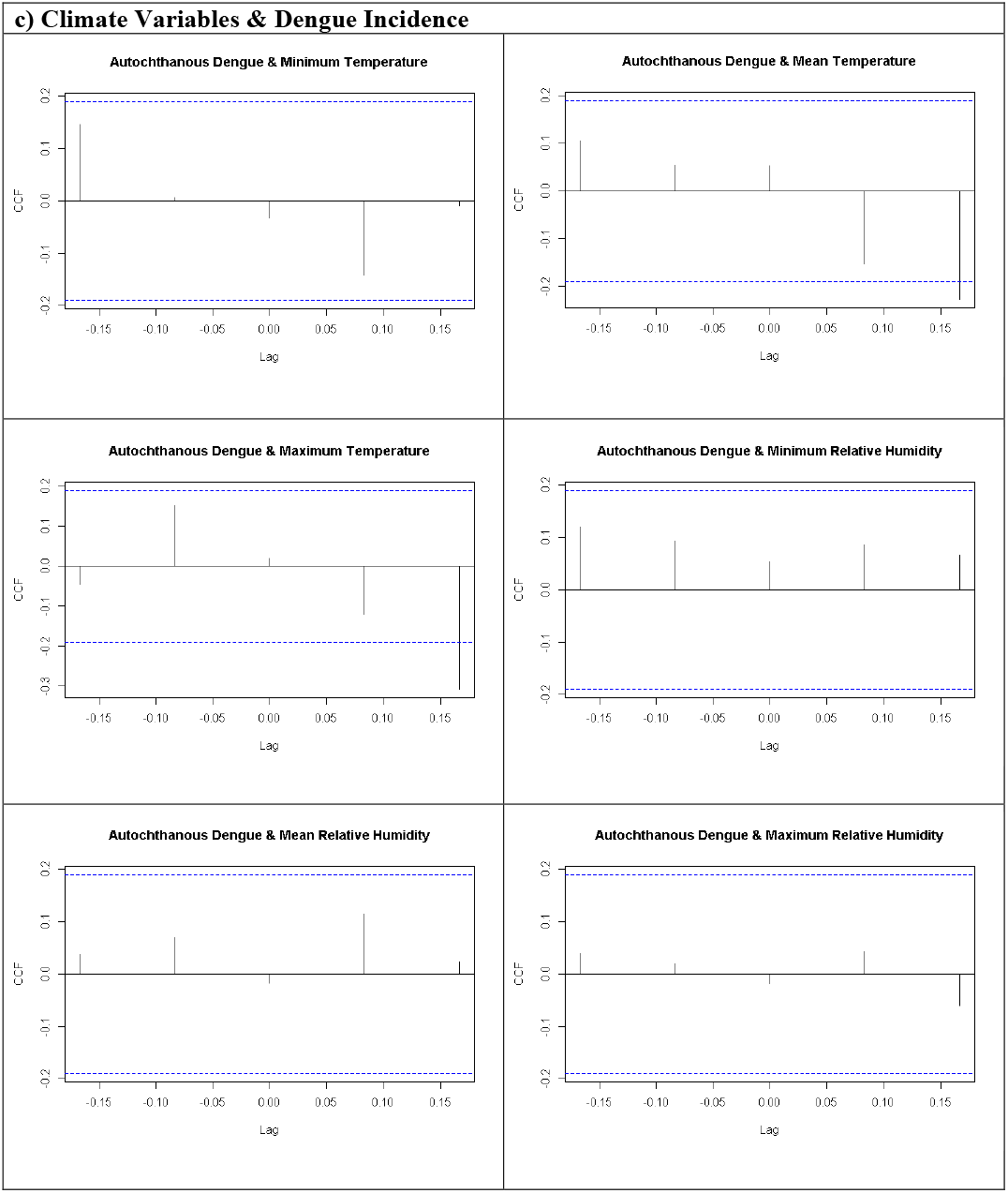
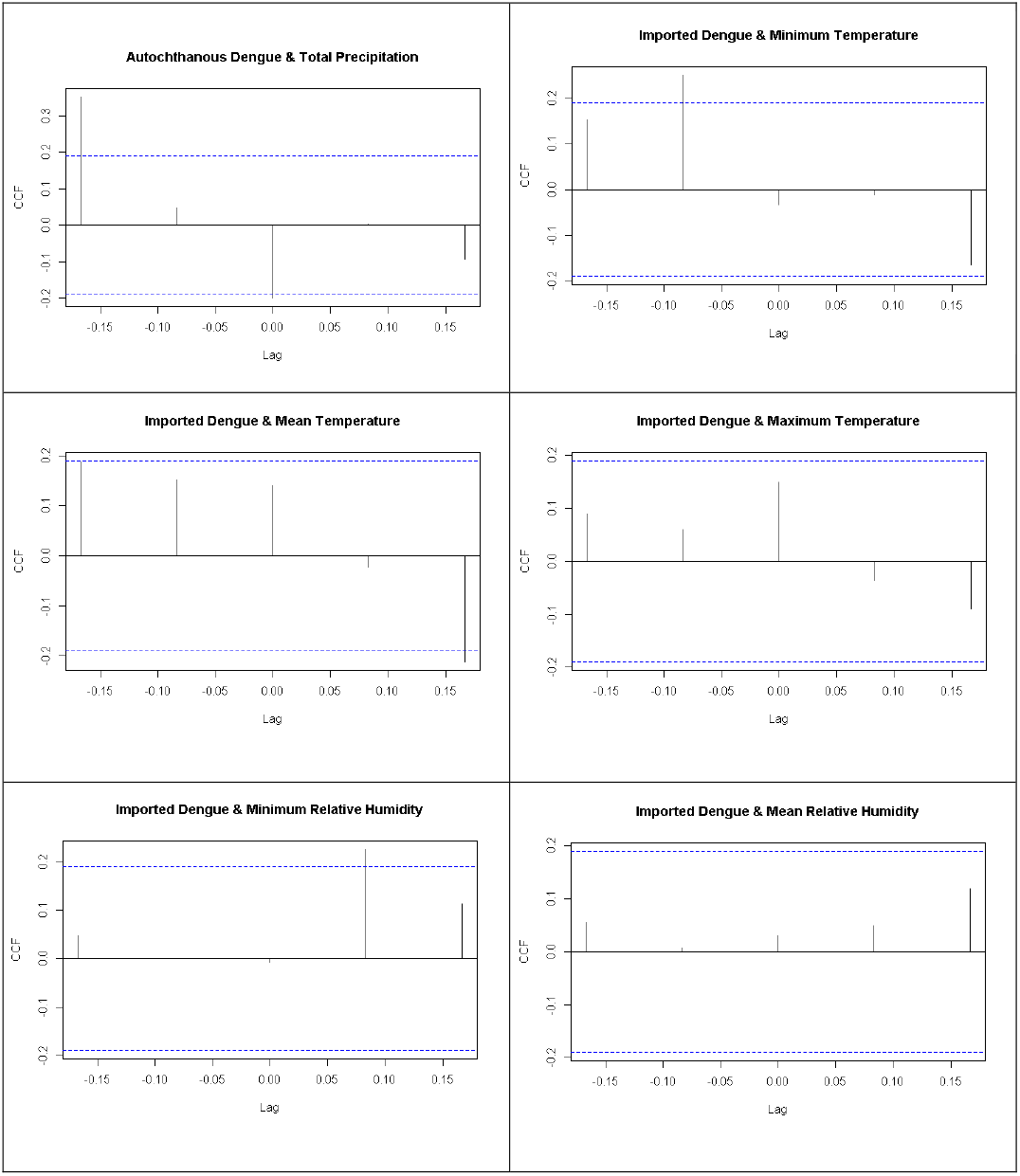
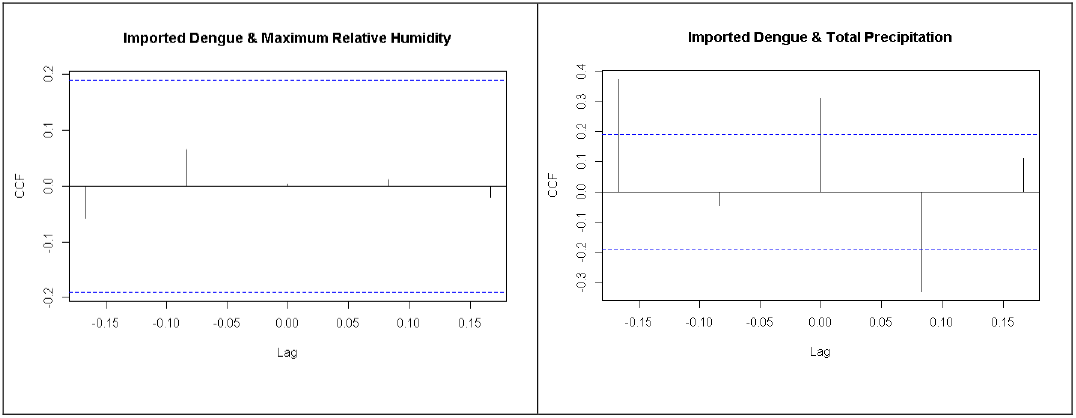
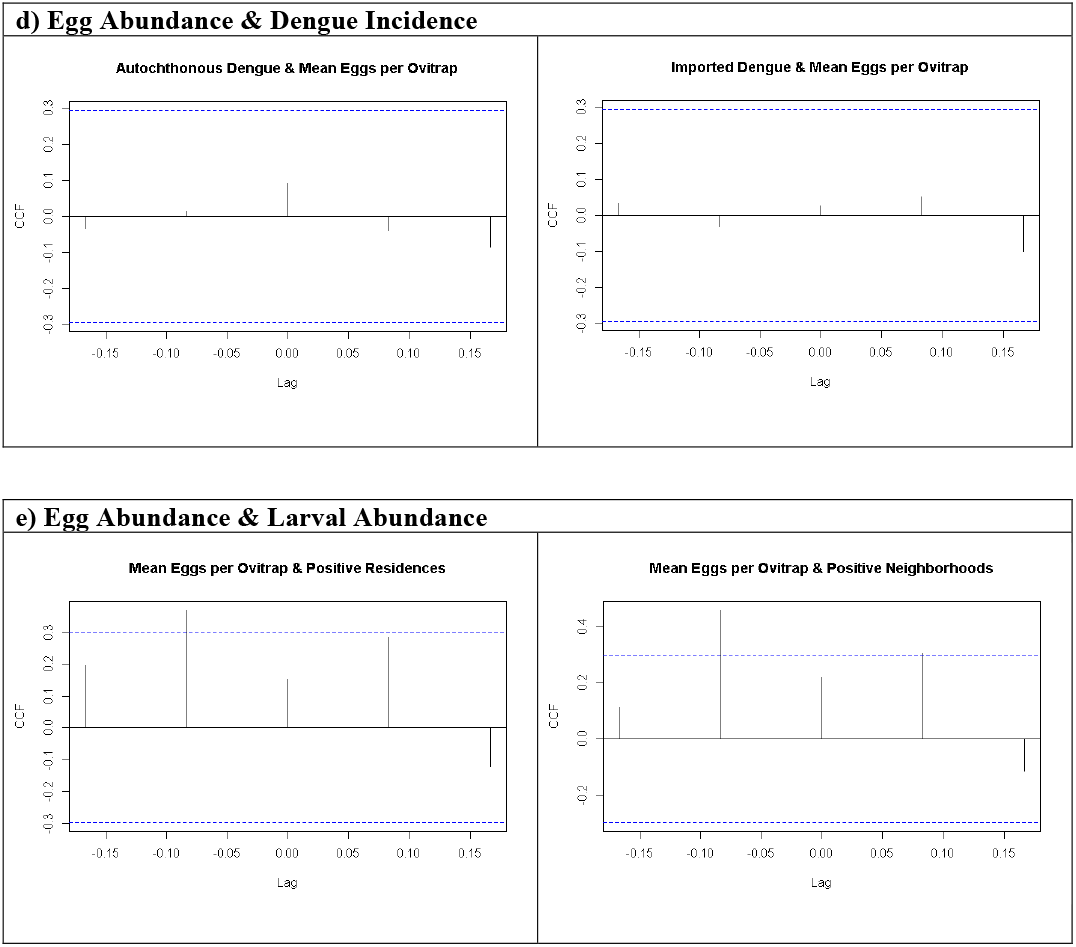
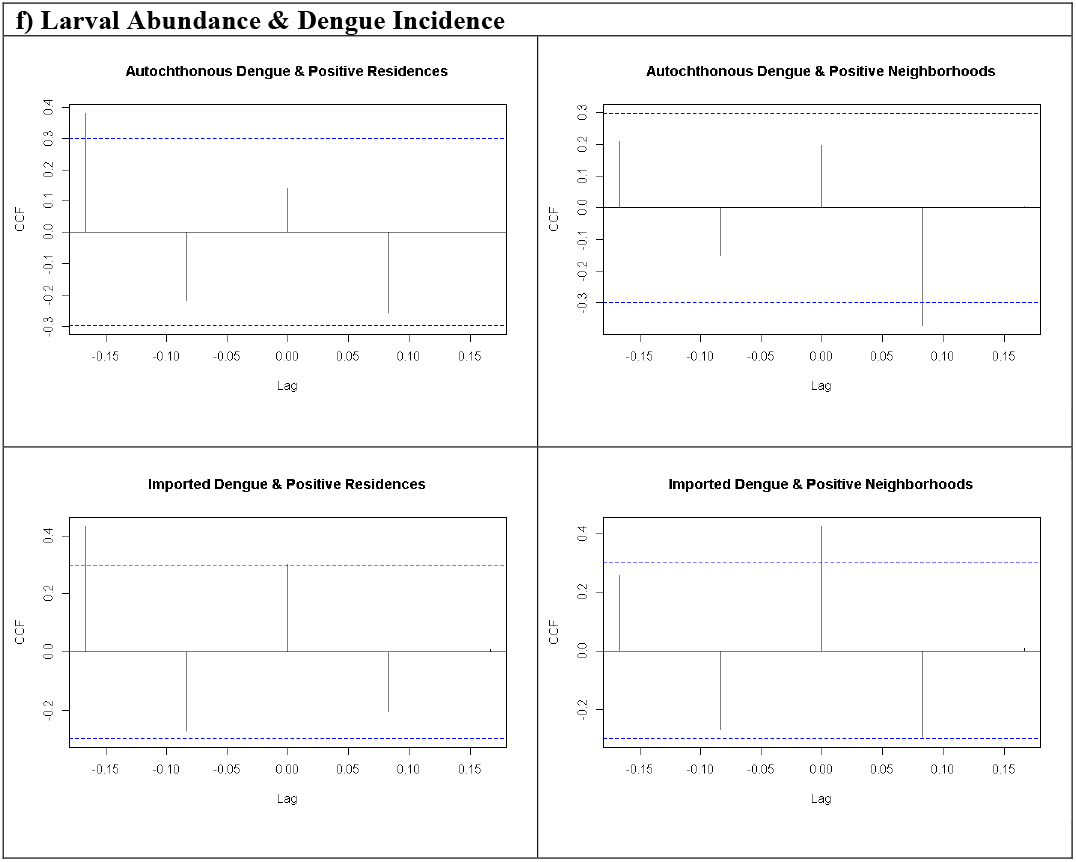

