## Supplementary figures and images for "A decade of arbovirus emergence in the temperate southern cone of South America: dengue, *Aedes aegypti* and climate dynamics in Córdoba, Argentina"

### Time series of monthly dengue, Aedes aegypti larval and Ae. aegypti eggs abundance.

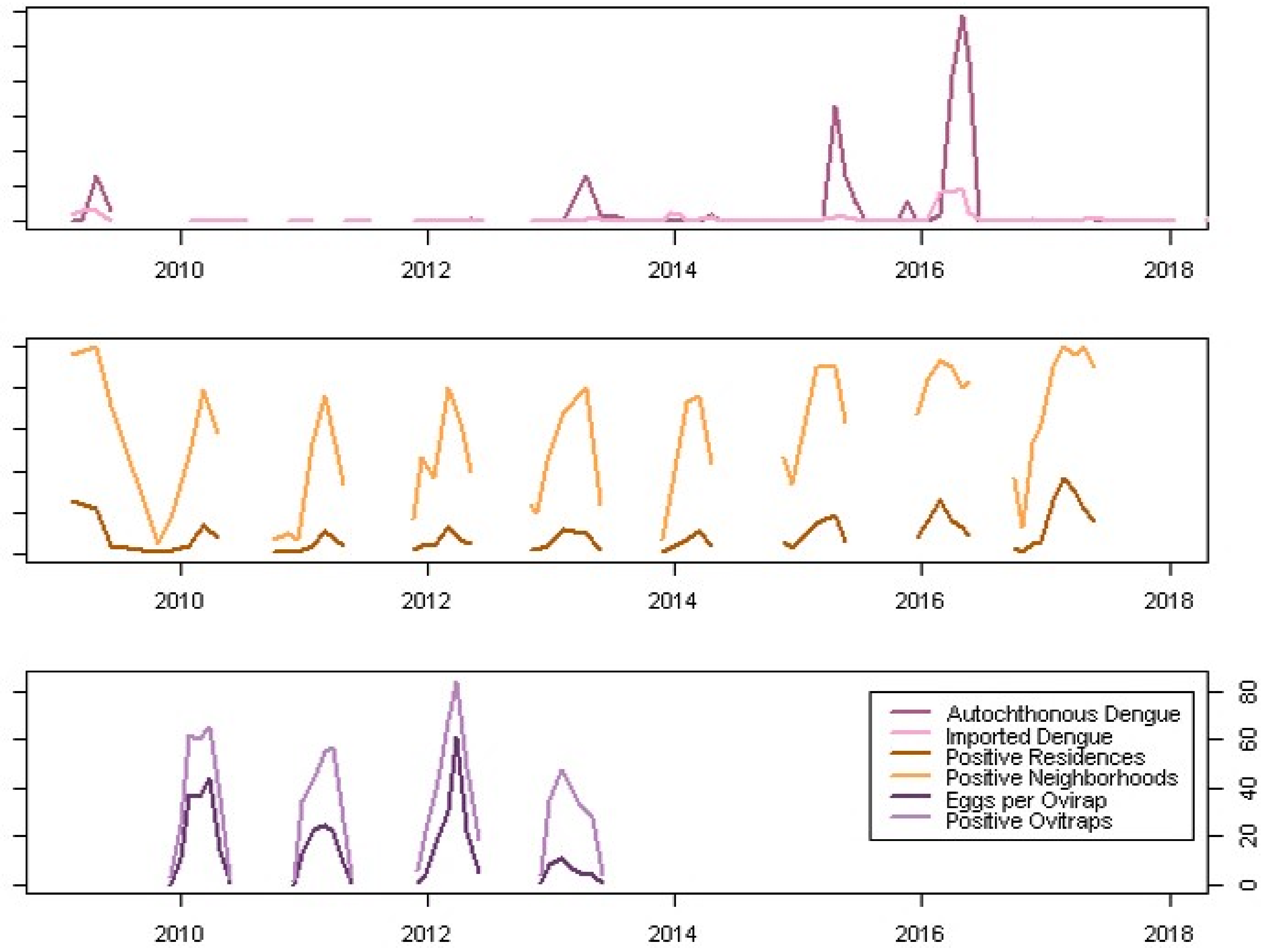

### Times series of monthly climate variables.

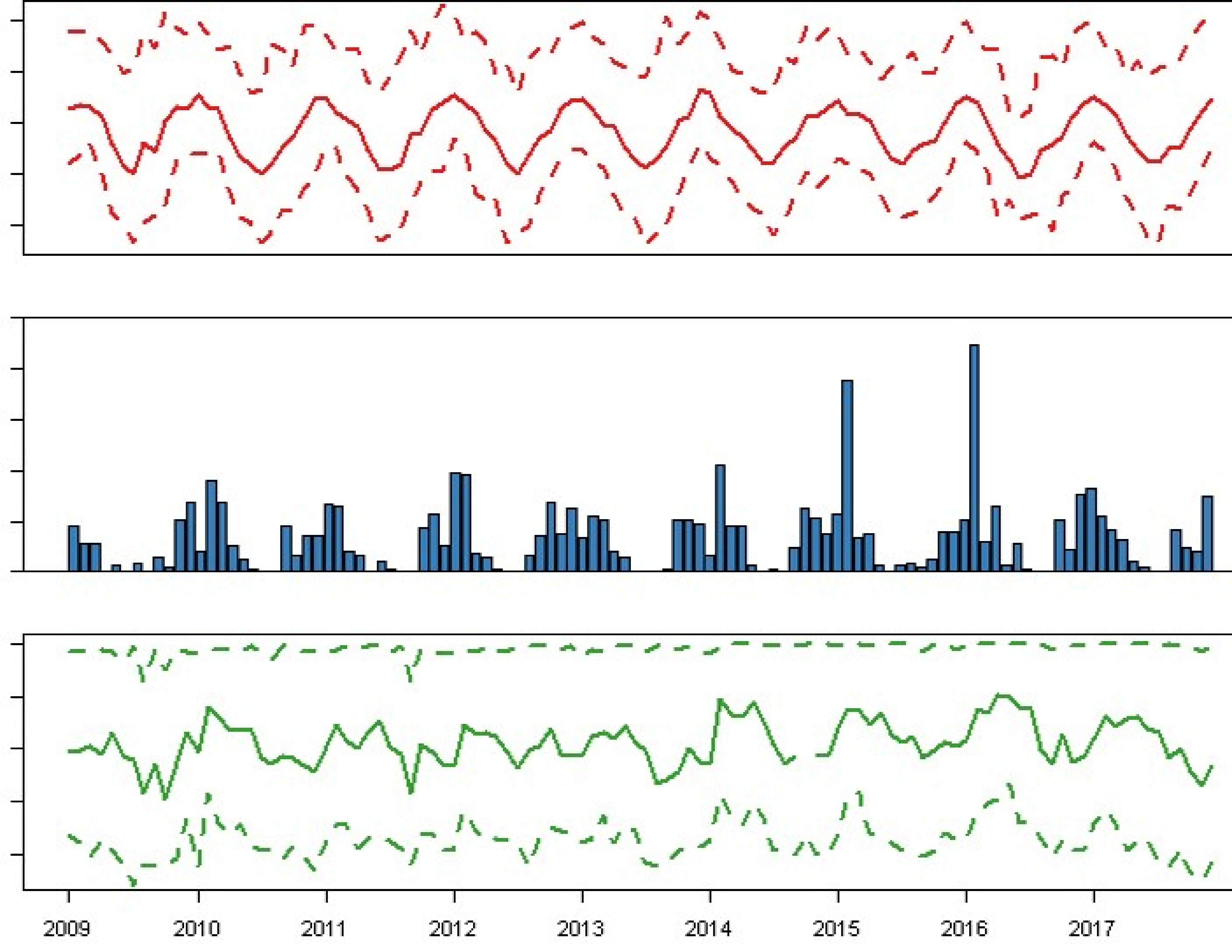
